# The foraging perspective on economic choice

**DOI:** 10.1101/190991

**Authors:** Benjamin Y. Hayden

## Abstract

Foraging theory offers an alternative foundation for understanding economic choice, one that sees economic choices as the outcome of psychological processes that evolved to help our ancestors search for food. Most of the choices encountered by foragers are between pursuing an encountered prey (accept) or ignoring it in favor of continued search (reject). Binary choices, which typically occur between simultaneously presented items, are special case, and are resolved through paired alternating accept-reject decisions limited by the narrow focus of attention. The foraging approach also holds out promise for helping to understand self-control and invites a reconceptualization of the mechanisms of binary choice, the relationship between choosing and stopping, and of the meaning of reward value.

**Highlights:**

- Foraging provides a basis for modeling economic choice based on adaptiveness
- Foraging choices are accept-reject; foraging models interpret binary choice accordingly
- The foraging view offers a different perspective on self-control decisions
- Economic and stopping decisions may have a common basis

## Introduction

The need to pursue and consume food is a core part of any animal’s behavioral repertoire and is a major driver of evolution. Decision-making is a key element of foraging - foragers must make tradeoffs that will maximize intake of desired prey and avoid unnecessary delay, risk, and effort. Foraging provides an approach to the psychology and neuroscience of economic decision-making, one that comes with a firm foundation, the principles of evolutionary adaptiveness (Pearson et al., 2014; Passingham and Wise, 2012). In this brief review, we provide a summary of the psychology and neuroscience of economic decision-making as seen through the lens of foraging.

## Foraging decisions are accept-reject decisions

For most foragers, the distribution of prey in the environment is patchy in space and ephemeral in time (Stephens and Krebs, 1986). As foragers search, prey are generally encountered one at a time and the forager’s decision is whether to pursue (accept) or ignore (reject) the prey item. This principle is true for both for a forager searching for individual prey organisms within a patch and foragers surveying multiple patches. The elemental decision in foraging, then, is the accept-reject decision and not, as in microeconomics, a binary choice between simultaneously presented items.

Although accept-reject decisions are ostensibly binary choices between two simultaneously appearing items (i.e. accept and reject), they function differently in practice. Pursuing a prey item is active, it leads to reward consumption (formally, it is exploitative), and it involves a change from the current state; capture normally triggers monitoring, adjustment, and learning processes (as does failed capture following an accept decision). Rejecting a prey item is passive, maintains the status quo (that is, continuing to search), and, during the subsequent search, returns the decision-maker to outwardly-oriented searching (formally, exploratory) mode.

## Insights into self-control from the accept/reject framework

One important example of the way the accept/reject framing matters is the performance of animal decision-makers in intertemporal choice (a.k.a. delay discounting) tasks. Such tasks, in which animals choose between delayed large rewards and immediate small rewards, are a mainstay of the psychology and neuroscience of self-control (reviewed in Hayden 2015). Animals generally appear impulsive, meaning they prefer a reward offered sooner even if it is less profitable. This observation is difficult to reconcile with evolutionary theory because it is highly maladaptive in the long run. However, intertemporal choice tasks, structured as binary choice tasks, will lead a decision-maker evolved to making accept-reject decisions to misattribute delays appear impulsive (Stephens et al., 2004; Bateson and Kacelnik, 1996). Indeed, in foraging tasks with a time component, ostensibly impulsive animals are almost perfectly patient (Blanchard and Hayden, 2015; Stephens and Anderson, 2001; Pearson et al., 2010; Carter and Redish, 2016).

Another important feature of foraging decisions absent from standard economic ones is the requirement for persistence in pursuing or handling a prey item after the decision itself it made. The neural mechanisms of persistence are just beginning to be understood (Hillman and Bilkey, 2013; McGuire and Kable, 2015; Wittmann, 2016; Rudebeck, 2006). One key ingredient in many persistence decisions is the need to maintain an ongoing representation of the value of the prey and to update that value continuously as the receipt of the prey gets closer in time (McGuire and Kable, 2015; Blanchard et al., 2015). Failures of this value updating process may help to explain failures of self-control, and treatments that modify this representation may help improve self-control.

## How accept-reject decisions are implemented

The key decision variable in accept-reject decision is profitability: the gain weighed against the cost of the item, including opportunity costs (Stephens and Krebs, 1986). Profitability is compared to a threshold, the average value of the environment (the analogous term in baseball statistics is the Mendoza Line). The most straightforward way to implement an accept-reject decision is to maintain a (dynamic) representation of the profitability of the foreground option and a (stable) representation of the profitability of the background and to compare them (Kolling et al., 2012; Wittmann et al., 2016; Hayden et al., 2011). Control systems in the brain then can modulate these representations regulate the threshold for accepting a presented option (Kolling et al, 2014; Hayden et al., 2011; Steiner and Redish, 2015).

Once an item is attended, accepting the option may be a type of default action; rejecting it would then be an alternative. If so, this framing would introduce an asymmetry into binary choice. That asymmetry should be visible in the brain, and indeed it is: the two option types are associated with activation in the ventromedial prefrontal cortex (vmPFC) and dorsal anterior cingulate cortex (dACC), for default and alternative, respectively (Boorman et al., 2013; Kolling et al., 2012). Presumably then choice is determined by competition processes between these two systems. The role of dACC in encoding the value of the alternative is also consistent with recordings of single neurons there, which show encoding of the rejected value on reject trials and of the delay - which corresponds to the opportunity cost of the accept decision - on accept trials (Blanchard and Hayden, 2014). And in a patch-leaving task, in which the decision to reject an option builds over several trials, responses of dACC neurons gradually increase as the value of rejection rises (Hayden et al., 2011). These discoveries about dACC offer a synoptic account of dACC function that was not available from standard conflict and comparator models based on conventional (non-foraging) tasks (Kolling et al., 2016; Heilbronner and Hayden, 2016).

## Are ostensibly binary choices really paired accept-reject choices?

Binary choice is the very core of microeconomics and understanding its neural basis is a central goal of neuroeconomics. Given the importance of accept/reject decisions in foraging, however, some scholars have argued that the binary choice is at least somewhat unnatural and in some cases an artificial laboratory construct (Kacelnik et al., 2011; Stephens et al., 2004). A forager whose brain is evolved for single encounters may treat the binary choice as two simultaneous accept/reject decisions. Key evidence for this decision mechanism comes from measures of reaction times and choice probabilities (Shapiro et al, 2008; Freidin et al, 2009; Vasconcelos et al, 2010; Pirrone et al, 2016). An implication of these results is that binary choice is better described as a paired race to threshold than a drift diffusion between two bounds.

Another psychological limitation on binary choice is the limited capacity of attention: we cannot bind abstract features (like value) to objects (offers) in the absence of attention, which is generally limited to a single spotlight (Treisman and Gelade, 1980). In a standard visual task with two spatially separate options, the spotlight of attention likely follows the locus of gaze or, sometimes, covert attention (Krajbich et al, 2011). In more abstract situations, such as when choosing between two possible options that are out of view (e.g. a monkey choosing which of two distant orchards to forage in), the locus of attention likely shifts in a more abstract manner, but still serially. Thus, it seems likely that when options are presented simultaneously, they are nonetheless evaluated and compared serially. Key evidence for this idea comes from a study that recorded ensemble activity in orbitofrontal cortex (OFC) in a simultaneous choice task (Rich and Wallis, 2016). Neuronal ensembles rapidly oscillated between two states corresponding to the two possible options, presumably tracking the focus of attention. Further evidence comes from the fact that, when attention is artificially controlled (by controlling gaze), ventromedial prefrontal cortex (vmPFC) and OFC preferentially track the values of attended offers (Lim et al, 2011, Strait et al, 2014; McGinty et al., 2016; Blanchard et al., 2015; Xie, Nie, and Yang, 2017).

## How can comparison occur in serial choice models?

If attention oscillates between single offers, and the brain signals the value of the attended offer only, how can a comparison occur? One possibility is that the brain computes the *relative*, not absolute, value of the attended offer (that is, the value difference or quotient). This relative value can be seen as a normalized representation of the value of the offer, but is sufficient to guide choice: if the difference is greater than zero, or if the quotient is greater than 1, the attended option can be selected. There is evidence that value representation in vmPFC is relative (Strait et al, 2014, Lim et al,, 2011) and may be relative in other areas as well (e.g. Hunt et al., 2014; Strait et al, 2015; Hunt et al, 2012; Klein-Flugge et al, 2016; Padoa-Schioppa, 2009). To implement choice, then, such normalized value representations must be subject to some downstream (or distributed) comparison-to-threshold process.

During serial shifts of attention, what is occurring during each epoch of sustained attention on one option? It seems likely that the brain is gradually accumulating evidence in favor of or against selecting that option (Krajbich et al, 2011; Pisauro et al, 2017). That evidence is presumably stochastic, because it reflects the output of multiple noisy channels. It seems likely that at least some of that sampling corresponds drawing recollections of stimulus and action value mappings from memory (Shadlen and Shohamy, 2016; Ludvig et al, 2014). These are then fed into one or more value buffers and compared to a threshold.

## Foraging suggests a unification of economic and stopping decisions

An accept/reject decision is a choice between actively changing the status quo or passively maintaining it and continuing to search; accepting involves performing a planned or primed motoric response; rejection involves withholding it. In other words, an accept/reject decision has much in common with a stopping decision. And binary choice, by extension, has much in common with a pair of interacting stopping decisions. This speculation, if true, is important because the neural mechanisms of stopping are relatively well understood, and applying this understanding to economic decisions could rapidly advance the neuroscience of economic choice (Logan et al., 2015; Aron et al, 2004; Hampshire and Sharp, 2015). Indeed, if economic choice ultimately boils down to stopping, there is an opportunity for a “grand unified theory” unifying the two types of decisions.

There is some tentative evidence that the neural circuitry involved in stopping is overlapping with the circuitry involved in economic decisions. The motor and premotor cortex, for example, have clearly defined roles in stopping decisions, and also have important and complementary roles in economic choices (Cisek and Kalaska, 2010; Cisek, 2012). More broadly, at least some evidence supports the idea that stopping is a distributed process that reflects activity of much of the prefrontal cortex (among other regions, Hampshire and Sharp, 2015); similar arguments have been made for economic choice (Cisek, 2012; Hunt and Hayden, 2017). In any case, future work on the relationship between stopping and choice is needed. Progress in this area promises to help shed light on important debates, such as how economic choice relates to selfcontrol (Berkman et al., 2017; Shenhav, 2017).

## Value as tentative commitment to a decision

Value is a construct that is convenient in economic models, but may not be explicitly computed; evidence that it is realized in the brain is equivocal (Hunt and Hayden, 2017; Vlaev et al., 2011). The brain has not evolved to compute value and then use that to drive choice; it has evolved to drive choice (Cisek and Kalaska, 2010). Indeed, a reasonable null hypothesis would be that the brain, as an evolved system, performs a gradual rotation from an input to an output space without a special amodal value representation in a middle layer. Such a rotation would lead to sensorimotor information - that is, the details of the positions of offers and actions leading to them, in ostensibly motor areas. Recent evidence supports the idea that such signals are indeed observed throughout the reward system (Strait et al, 2016; McGinty et al., 2016; Bryden and Roesch, 2015; Luk and Wallis, 2013; Cisek, 2012). What we call value, then, may really a tentative commitment to accepting an offer, or, more abstractly, to a proposition (cf. Shadlen et al, 2008). In early layers, that proposition may be toward identifying the stimulus, in middle layers, it may be towards signals that can influence the action or the goal, and in later layers it may be towards the action associated with choosing it (Cisek, 2012; Hunt and Hayden, 2017; Hunt et al, 2014; Cisek and Kalaska, 2010). Future studies will be necessary to test this idea; such studies are most likely to be informative if they are ethologically relevant, that is, if they embed the decision-maker in as natural an environment as possible (Pearson et al., 2014).

---

**Figure 1.**
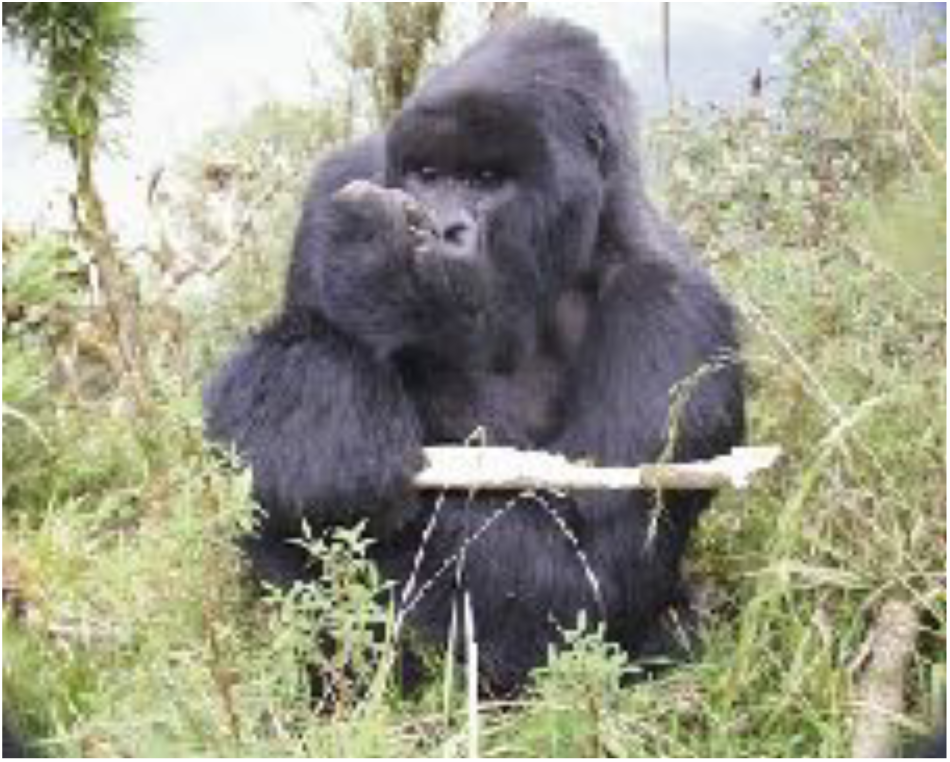
Animals in natural environments, such as this gorilla at the Karisoke Research Center in Rwanda, generally encounter prey one at a time. Their decision-making strategies are molded by those encounters and are centered on accept-reject decisions. (Photo credit: Jessica Cantlon).

**Figure 2.**
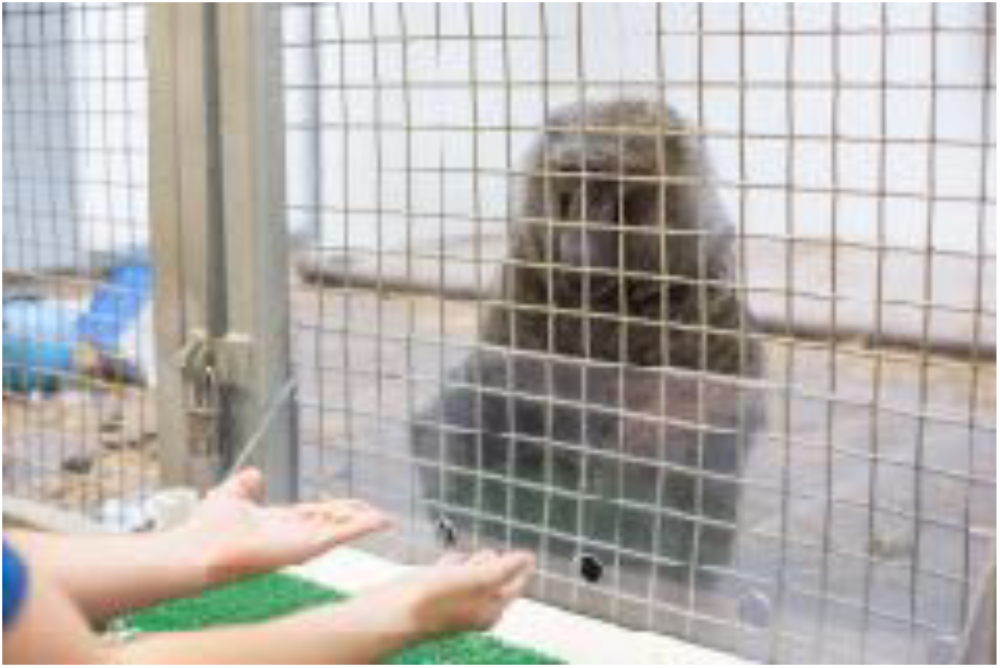
Animals in the laboratory, such as this baboon at the Seneca Park Zoo, are often faced with binary decisions. Some research suggests that those decisions are made as interleaved accept-reject decisions of each option. (Photo credit: Jessica Cantlon).

